# Diet changes thermal acclimation capacity, but not acclimation rate in a marine ectotherm (*Girella nigricans*) during warming

**DOI:** 10.1101/2022.10.25.513746

**Authors:** Emily A. Hardison, Gail D. Schwieterman, Erika J. Eliason

## Abstract

Global climate change is increasing thermal variability in coastal marine environments and the frequency, intensity, and duration of marine heatwaves. At the same time, nutritional resources are being altered by anthropogenic environmental changes. Marine ectotherms often cope with changes in temperature through physiological acclimation, which can take several weeks to occur and is a nutritionally demanding process. Here, we tested the hypothesis that different ecologically relevant diets (omnivorous, herbivorous, carnivorous) can impact thermal acclimation rate and capacity, using a temperate omnivorous fish as a model (opaleye; *Girella nigricans)*. We measured acute thermal performance curves for maximum heart rate because cardiac function has been observed to set upper thermal limits in ectotherms. Opaleye acclimated rapidly after warming, but their thermal limits and acclimation rate were not affected by diet. However, the fish’s acclimation capacity for maximum heart rate was sensitive to diet, with fish in the herbivorous treatment displaying the smallest change in heart rate throughout acclimation. Mechanistically, ventricle fatty acid composition differed with diet treatment and was significantly related to cardiac performance in ways consistent with homoviscous adaptation. Our results suggest that diet is an important, but often overlooked, determinant of thermal performance in ectotherms on environmentally relevant timescales.

## Introduction

Understanding the mechanisms driving thermal tolerance in ectotherms is a primary focus in ecological physiology. Temperature is one of the most important environmental factors governing the physiology, behavior, and ecology of ectotherms [1,2]. Temperature profiles vary substantially across environments, and thus endemic ectotherms are adapted to tolerate different thermal regimes [3]. Temperate environments are especially thermally dynamic, with changes in temperature occurring on acute (diurnal), intermediate (seasonal upwelling and marine heatwaves), and long term (annual – decadal) timescales [4–6]. Importantly, climate change is expected to increase the severity and duration of these natural (i.e., upwelling, diurnal cycles) and extreme (i.e., heat waves) sources of temperature variation [6,7]. To survive in these dynamic environments, ectotherms must be able to cope with changes in environmental temperature across all relevant timescales [6,8,9]. As such, animals inhabiting these variable environments generally possess the capacity for high levels of thermal plasticity, otherwise known as thermal acclimation capacity.

Thermal acclimation occurs through a series of phenotypic changes across levels of biological organization that optimize biological rates, minimize waste production or cellular damage, or conserve energy [10–13]. Importantly, each phenotypic change takes time. Acclimating quickly may be just as, if not more important than full acclimation capacity [9,14–16], especially in the context of an increasingly variable environment. As the heart is essential for transporting nutrients, waste, immune cells, oxygen, etc. around the body, it is thought to govern thermal tolerance in several ectotherms [17] and has been shown to rapidly acclimate to high temperatures in some temperate fish [13,18]. Given that acclimation is a remodeling process, successful and efficient acclimation requires energy and nutrients. Thus, the rate of acclimation and overall acclimation capacity for cardiac thermal tolerance may be dependent on the quality and quantity of what an ectotherm has eaten.

Climate change is predicted to alter food availability, nutritional quality, and diet preference for several ectotherms [19–22], yet the interaction of diet and temperature on thermal performance is understudied. Animals have a remarkable capacity to vary their diet to meet their nutritional requirements through changes in their consumption rate or diet selection [23–26]. Generalist ectotherms, like omnivores, can choose between foraging for plants or animals to meet their nutritional needs. Interestingly, some omnivorous aquatic ectotherms vary the ratio of plant to animal in their diet with temperature, suggesting that the nutritional needs of these ectotherms are temperature dependent [21]. The physiological consequences of these diet shifts are not well understood but may be associated with altered performance and thermal tolerance [27–29]. An outstanding question is whether the ratio of plant to animal in an omnivore’s diet affects their thermal plasticity. More specifically, can diet affect thermal acclimation rates and capacity, and if so, what mechanisms underly these differences.

Diet quality varies dramatically across prey types. For omnivores, plants tend to be higher in mineral content, antioxidants, and complex carbohydrates, while animal diets tend to have higher protein and lipid content. In general, fatty acid (FA) composition also differs between plants, algae, and animals, although the exact composition is also dependent on the environment (i.e., terrestrial, marine, freshwater) and conditions (e.g., environmental oxygen, temperature) [30]. For example, the global availability of essential omega-3 fatty acids, docosahexaenoic acid (DHA) and eicosapentaenoic acid (EPA), are expected to decline due to increasing temperatures associated with climate change [31]. FA composition can influence membrane fluidity, thermal limits [32–33], cardiac performance [34], swimming performance [35], growth [36], cognitive function [37], and diet preference [29], suggesting that variation in FA composition may be a primary mechanism underlying thermal performance differences in ectotherms that change their diet with temperature.

The objective of this study was to determine whether diet impacts 1) the rate and 2) the capacity of cardiac thermal acclimation in a temperate omnivorous fish, opaleye (*Girella nigricans*), during a simulated warming event. We measured thermal performance of maximum heart rate (*f*_Hmax_) as it is thought to limit thermal tolerance in fishes, and heart rate is governed by processes occurring across biological lipid membranes [17]. Opaleye were fed either a fully herbivorous (ad lib red algae, *Gracilaria pacifica*), carnivorous (ad lib brine shrimp, *Artemia* sp.), or omnivorous (free choice between algae and *Artemia* sp.) diet for two weeks at 12°C. We then increased temperature to 20°C, simulating an acute marine warming event and periodically measured cardiac thermal performance over the next two weeks while the fish were acclimating to 20°C. We predicted that the lower energy, lipid, and protein content in the algae compared to *Artemia* sp. would result in lower acclimation capacity, slower acclimation rates, and reduced upper thermal limits compared to the carnivorous treatment. Further, that fish in the omnivorous treatment would have the highest performance due to the greater diversity of nutrients available, high energetic content of the diet, and flexibility to choose between the algae and *Artemia* sp. To examine the mechanisms underlying observed differences in cardiac performance, we measured the FA composition of the fish’s ventricles and related FA composition to heart rate and thermal tolerance.

## Methods

### Fish Collection

Juvenile opaleye were collected in May-August, 2021 by hook and line from Santa Barbara Harbor, California, USA (34.40829, −119.691389). Fish were transported in coolers (>70% air saturation) to the University of California, Santa Barbara and held in 95 L fiberglass flow-through seawater tanks (14 fish per tank, 4x turnover h^−1^). Prior to the start of acclimation, fish were held at ambient conditions (ranged ~12.5-17.5°C throughout the study) for <3 weeks and fed *ad libitum* omnivorous diets (*Gracilaria pacifica* from the cultured abalone farm in Goleta, CA and *Artemia* sp. from brineshrijmpdirect.com). All protocols were approved by the Institutional Animal Care and Use Committee at the University of California, Santa Barbara.

### Acclimation and Diet Treatments

Opaleye were held at 12°C and fed *ad libitum* one of the three diet treatments for two weeks (4-5 tank replicates per treatment). The diet treatments were 1) carnivorous (*Artemia* sp.), 2) herbivorous (red macro algae native to California called *Gracilaria pacifica*), and 3) choice omnivorous (*Artemia* sp. and algae). Consistent with other aquatic omnivorous ectotherms, opaleye increase algae relative to prey consumption in the warmer part of their geographic range [38]. After two weeks, fish were tested using a cardiac thermal tolerance test called an Arrhenius breakpoint temperature test (ABT test; outlined below). One to two days later, temperature in the holding tanks was raised over ~10-12 h from 12 to 20°C (representative of the low and high seasonal temperatures experienced in Santa Barbara, CA). As temperate intertidal fish, this rate of temperature change is high for Santa Barbara, however not unheard of given the ecology of the species and a common rate of change in more Southern parts of their range. Following the temperature change, individuals from each treatment were tested using an independent sampling design on day 1 (post-change), 3, 7, and 14-15 for their upper cardiac thermal limits using the same ABT test. The experiment was replicated 4 times to ensure adequate sample sizes for each timepoint. A 5^th^ replicate was also conducted until day 3 post temperature change to increase the sample sizes at earlier timepoints. We also included one tank of fish that were acclimated to 20°C for 4 weeks and tested on week 2 and 4 of acclimation (Figure S1).

Each tank started with 14 fish per tank. During each sampling day, 2-4 fish from each treatment were removed and tested. Temperature and dissolved oxygen content were monitored daily by hand using an Omega Thermocouple (Omega Engineering INC, Norwalk, CT, USA) and an Oxyguard handy Polaris 2 (OxyGuard International A/S, Farum, Denmark). Oxygen was maintained at >80% air saturation throughout the study. Average temperature per treatment was 12.2 ± 0.5°C and 20.0 ± 0.6°C across replicates (mean ± SD; determined from in tank Thermochron 4K iButtons programmed to record every 20 minutes; accuracy ± 1°C and resolution to 0.5°C). Replicate level temperature data is provided in Table S1. Fish were held under a 14:10 h light: dark cycle.

### Direct Field Sampling

Juvenile opaleye were collected in the August of 2021 (avg temp for 2-weeks prior was 18.4 ± 1.0°C; mean ± SD) from Santa Barbara Harbor using hook and line. ≤ 4 fish were caught per day for 3 days. Fish were immediately transported back to the University of California, Santa Barbara and an Arrhenius breakpoint temperature test was conducted within 3 h of being caught using water from the harbor. This was done to examine how our results compared to seasonally warm-acclimatized opaleye in the wild.

### Dissections and Stomach Contents

At the end of each test, liver and ventricle from the fish were weighed, and the ventricle was flash frozen for later analysis. Fish were not fasted before testing to allow for collection of stomach contents. At the time of sampling, stomach contents were weighed and sorted in the omnivorous treatment to estimate the proportion of algae to brine shrimp and whether algae had been detected in the stomach contents. Omnivorous diet information is provided in Table S2.

### Proximate Analysis

Frozen fish remains on day 14 (after 2 weeks at 20°C and 4 weeks on the treatment diet) were homogenized using a Fisher Brand Bead Mill 24 and subsamples of the homogenate were weighed and freeze dried (Labconco Lyophilizer). Subsamples of algae and *Artemia* sp. were also freeze dried for proximate analysis (see Table S3). Protein, lipid and ash content were estimated as described in Hardison *et al*. [28]. Total fatty acid composition was determined on ventricles (n = 9 per treatment and timepoint) and diet samples (n = 6 per diet) by gas chromatography/mass spectroscopy (GC-MS) at the University of California, San Diego Lipidomics Core using methods outlined in Quehenberger *et al*. [39] (see Table S4, Table S5). In calculating measures of double bond index (DBI) and chain length, fatty acids that were “not detected” in the sample were assumed to be 0.

### Thermal Limits: Arrhenius Breakpoint Test

ABT tests on the heart were conducted as outlined in Hardison *et al*. [28] and Schwieterman *et al*. [40] (N = 7-15 per timepoint and per diet treatment; see Table S6 for specific values). Briefly, fish were anesthetized in seawater containing 80 mg L^−1^ buffered MS-222 and then placed in an experimental sling in a 10 L test tank which contained a maintenance dose of buffered 65mg L^−1^ MS-222. Water was circulated continuously past the gills throughout the test and stainless-steel needle tip electrodes (ADInstruments INC, Colorado Springs, CO, USA) were shallowly inserted under the skin to detect an ECG signal. The signal was amplified using a Dual Bio Amp (ADInstruments INC, Colorado Springs, CO, USA) and filtered (Filters: 60 hz Notch filter; Mains filter; Low-Pass: 1 Kz; High Pass: 10 hz; Range: 1-2 mV).

After a 15 min equilibration period at the acclimation temperature, atropine sulfate was injected intraperitonially (1.2 mg kg^−1^ in 0.9% NaCl) to block vagal tone. 15 minutes later, isoproterenol was injected intraperitonially (4 ug kg^−1^ in 0.9% NaCl) to maximally stimulate ß-adrenoreceptors. In 20°C acclimated fish, water temperature was cooled from 20 to 18°C before the start of the test. This was done to provide additional *f*_Hmax_ data for breakpoint temperature calculations.

After isoproterenol injection, fish were given a 30-minute equilibration period. Then water temperature was heated (warm ABT test) at 10°C h^−1^ (Polystat recirculating heater/chiller; Cole-Palmer, Vernon Hills, IL, USA) while continuously recording an ECG trace. At each 1°C interval, *f*_Hmax_ and temperature were stabilized for 30 seconds to record an average value for heart rate at the temperature. This procedure was repeated until the onset of cardiac arrythmia (T_ARR_). The fish were kept in the test until the next 1°C interval after T_ARR_ (<6 min) to ensure that the decrease in heart rate following cardiac arrythmia was captured for fitting thermal performance curves. Fish were omitted from the analysis when experimental error occurred (e.g., water pump failure, drug injection complications, or ECG signal was too noisy for software to interpret; 11% of individuals tested). Cold ABT tests were only performed on the last day of testing (i.e., day 14/15 post temperature switch; N = 5-7 per diet). The same rate of temperature change (decrease by 10°C h^−1^) was targeted; however, below 11°C, the rate of decrease in temperature was harder to maintain and deviated occasionally from the desired rate (~0.3-1 °C every 6 min). We do not anticipate that this impacted the results, as Casselman *et al*. [41] found that slower rates of temperature change did not impact the ABT test results. All fish were immediately euthanized and dissected at the end of the test.

### Data Analysis for Arrhenius Breakpoint Test

At each 1°C temperature increment, *f*_Hmax_ was calculated from 15 continuous seconds of ECG recordings using automated ECG analysis software in LabChart (AD Instruments; www.adinstruments.com)[28]. Three thermal limits were calculated from the warm ABT test (T_AB_, T_PEAK_, T_ARR_). The lowest thermal limit (T_AB_) represents when the heart first starts showing signs of impairment due to temperature. For example, T_AB_ was comparable to the optimal temperature window for aerobic scope in Pacific salmon [41], rainbow trout [42], and goldfish [43]. T_AB_ was defined as the temperature corresponding to a breakpoint in a plot of log(*f*_Hmax_) against the Arrhenius temperature. This breakpoint analysis was conducted using *segmented* package (v1.1-0)[44] in R. Note that T_AB_ may be artificially high in the early time points because the test started at a higher temperature (18 instead of 12°C). For the warm ABT test, the highest *f*_Hmax_ recorded during any 15 s measurement period was designated as the overall peak heart rate (Peak *f*_Hmax_). During the cold ABT test, the minimum heart rate (min *f*_Hmax_) was defined as the lowest *f*_Hmax_ recorded during any 15 s measurement period. The temperature in which peak *f*_Hmax_ occurred was defined as the peak temperature (T_PEAK_). T_PEAK_ indicates the temperature where any further increases in temperature are not met with corresponding increases in heart rate. This indicates an important transition temperature where cardiac arrythmia usually follows shortly thereafter. Finally, T_ARR_ indicates the temperature where the heart is no longer able to maintain rhythmic beating. At 12 and 20°C, T_ARR_ occurs at temperatures ~2.7-3.8°C lower than traditional measures of thermal tolerance in opaleye (i.e., critical thermal maxima; CT_max_) and thus, represents a more functional upper thermal limit for these fish [28].

### Statistical analysis

All data were statistically analyzed using R (version 3.5.1). All metrics were investigated for normality using Shapiro–Wilk tests and quantile–quantile plots, and for heteroscedasticity using Levene’s test. Acclimation rate data were statistically analyzed (significance level α=0.05) using a 2-way ANOVA where replicate number was included as a random effect (lme4 v1.1-19)[45] with post hoc Tukey’s tests for significant differences when significant main effects were detected. In all 2-way ANOVA tests, the interaction between diet and timepoint was tested for and excluded when non-significant. Polynomial curves were fitted to acclimation capacity *f*_Hmax_ data (i.e., data from fish held for 2 weeks at 12°C and 2 weeks at 20°C) and compared using Schwarz Information Criterion (SIC), where the fit with the lowest SIC score was assigned the best fit model, but all models with ΔSIC<7 were considered (Table S7) [46]. Replicate number and individual were included as random effects.

To simplify fatty acid analysis, the DBI and average chain length were calculated for each sample. 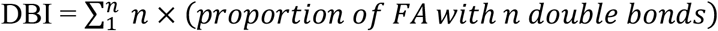, where n = the number of double bonds in a type of fatty acids. 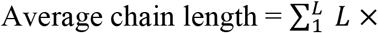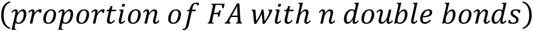, where L = the number of carbons in the fatty acid chain. Linear regressions between DBI or average chain length and all cardiac parameters (*f*_Hmax_, peak *f*_Hmax_, T_ARR_, T_PEAK_, T_AB_) were assessed. Residual plots of each linear regression model were visually assessed in R to investigate normality and heteroscedasticity. Complete dietary and tissue sample FA analysis (i.e., mean ± SEM for all FA’s measured) is provided in the supplement (Table S4, Table S5).

## Results

### Acclimation Capacity

The thermal acclimation capacity for *f*_Hmax_ was influenced by diet in opaleye (Figure 1, Table S8). Model selection revealed strong evidence of an interactive effect of diet and temperature on the thermal performance curve for *f*_Hmax_ in the warm ABT test (Table S7). The best fit model by SIC was a third-order polynomial curve that incorporated an interaction of acclimation temperature (12 or 20°C), diet (herbivorous, omnivorous, carnivorous), and acute test temperature (Table S7). The acute TPC for *f*_Hmax_ followed a traditional shape with a negative skew, where *f*_Hmax_ increased with temperature until T_AB_, at which point *f*_Hmax_ began leveling off until T_PEAK_. T_ARR_ generally occurred shortly after T_PEAK_.

**Figure. 1.**
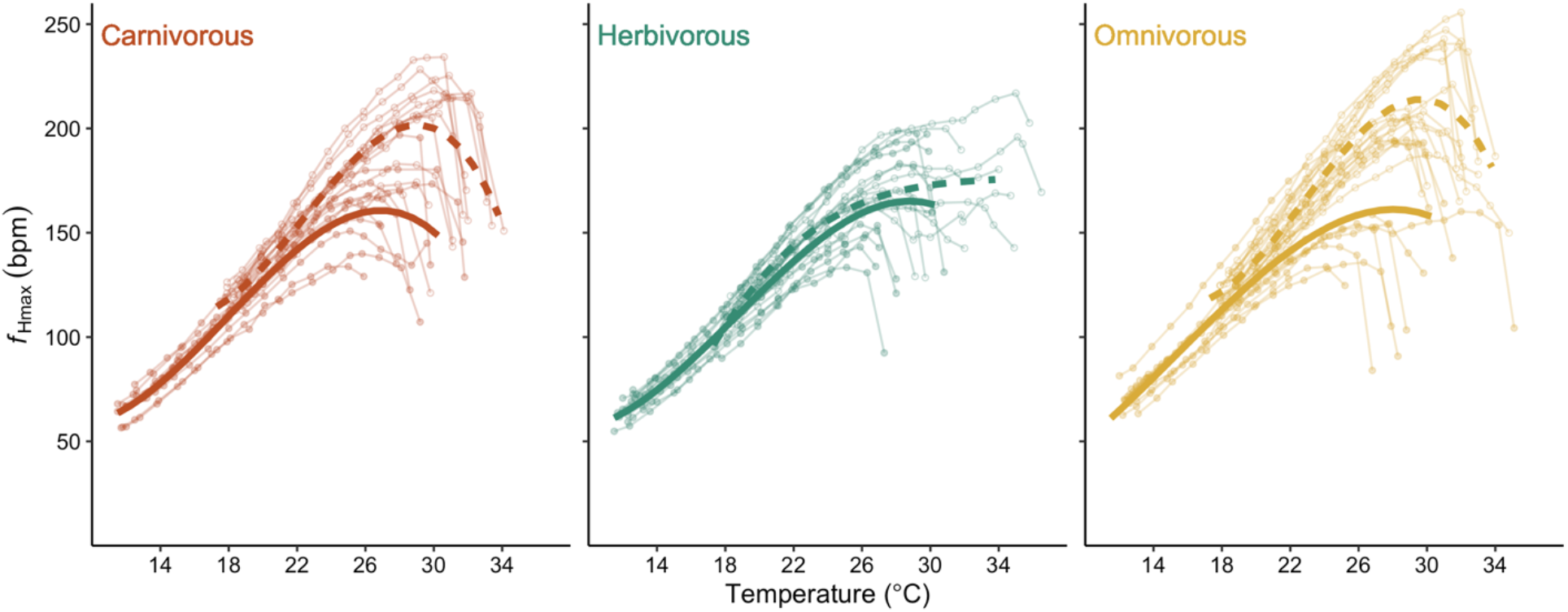
Dietary effects on cardiac thermal acclimation capacity. Individual- and treatment-level responses of max heart rate (*f*_Hmax_) of opaleye during acute warming. Panels show fish from different diet treatments (from left to right: carnivorous, herbivorous, omnivorous) that were tested after 2 weeks acclimation to 12°C (closed circles) and 20°C (open circles). Curves indicate treatment level effects after 12°C acclimation (solid line) and 20°C (dashed line) and are third-order polynomials that account for the interaction diet×timepoint×acute temperature with random effects of individual fish id and replicate number. This was determined to be the best fit model by BIC (Table S7).

Upper thermal limits (T_AB_, T_PEAK_, T_ARR_) increased by an average of 3.3-3.9°C (irrespective of diet) when switched from 12 to 20°C (Figure 2, Table S6). While diet did not affect any thermal limits (diet effect for T_AB_ p = 0.1647, T_PEAK_ p = 0.2269, T_ARR_ p = 0.1998, Table S6), it did impact the height of the acute TPC. For the omnivorous and carnivorous treatments, peak *f*_Hmax_ increased during acclimation from 12 to 20°C (Figure 3). In contrast, the herbivorous treatment did not alter peak *f*_Hmax_ in response to warm acclimation (p = 0.9856 from Tukey HSD comparing herbivore treatment at 12°C to the herbivore treatment after 2-weeks at 20°C). Accordingly, the herbivorous treatment had an overall Δ peak *f*_Hmax_ (peak *f*_Hmax at 20°C_ - peak *f*_Hmax at 12°C_) of 12.62 bpm compared to 55.10 and 43.96 bpm for the omnivorous and carnivorous treatments, respectively.

**Figure 2.**
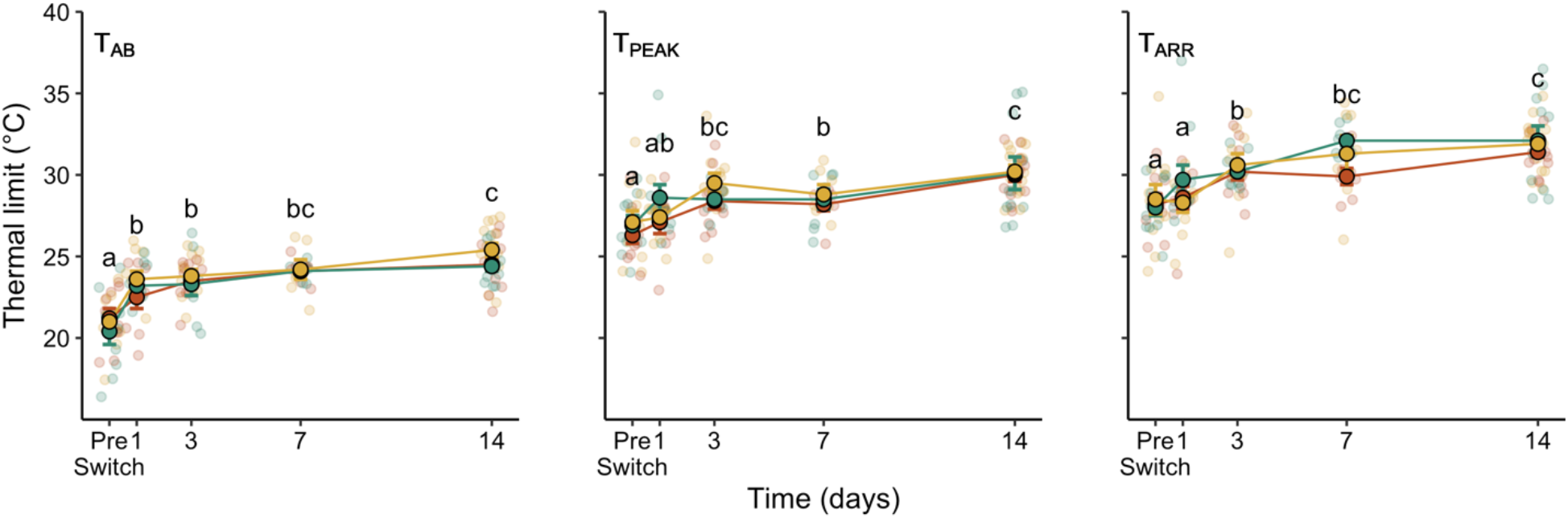
Dietary effects on thermal acclimation rate for cardiac thermal limits. Individual- and treatment-level acclimation rate responses for upper thermal limits (T_AB_, T_PEAK_, T_ARR_) across diet treatments and timepoints in opaleye. Panels and colors distinguish diet treatments (from left to right: carnivorous, herbivorous, omnivorous). Individual responses are indicated by the transluscent points, and treatment level mean (± SEM) are overlayed with connecting lines. Letters indicate significant differences among time points (p <0.05) by Tukey HSD. There were no significant effects of diet on any thermal limits.

**Figure 3.**
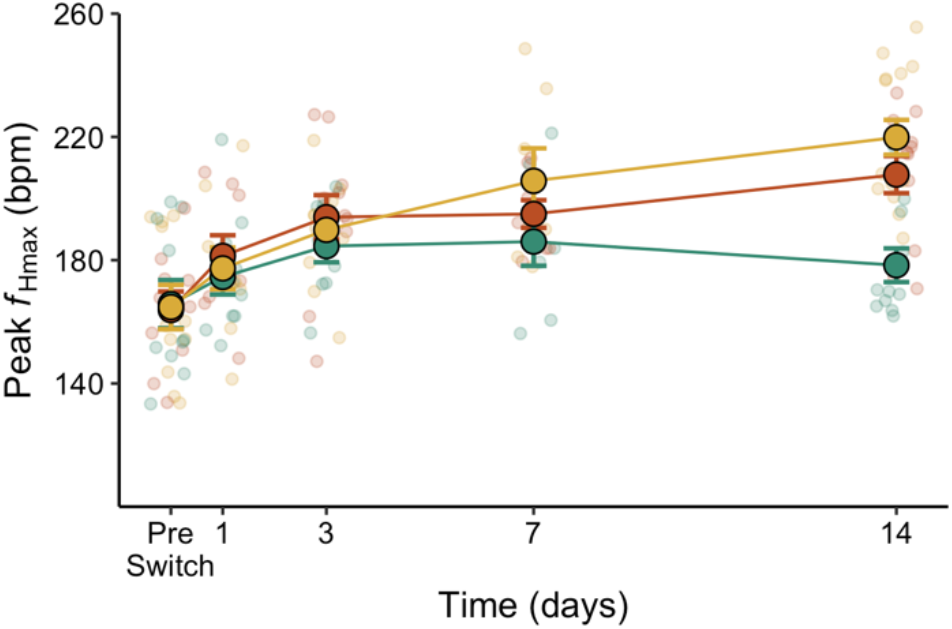
Dietary effects on thermal acclimation rate for peak *f*_Hmax_. Individual- and treatment-level acclimation rate responses in peak *f*_Hmax_ across diet treatments and timepoints in opaleye. Colors distinguish diet treatments (red: carnivorous, green: herbivorous, yellow: omnivorous). Individual responses are indicated by the transparent circles. Treatment level mean (± SEM) are overlayed with connecting lines. There was a significant interactive effect of diet and timepoint (P = 0.0128; Table S8) on peak *f*_Hmax_.

### Acclimation Rate

The majority of the acclimation response for thermal limits occurred in the first 3 days post temperature change, with an average of 69.2% of the response for T_AB_, 60.6% for T_PEAK_, and 60% for T_ARR_ occurring in the first 3 days at 20°C, irrespective of diet (Table S9). Diet did not significantly impact any thermal limit at any timepoint during acclimation from 12 to 20°C (Figure 2, Table S8). However, there was an interactive effect of diet and timepoint on peak *f*_Hmax_, where it was lower in the herbivorous treatment compared to the omnivorous (23.3% higher) and carnivorous (16.5% higher) treatments after the 2-week acclimation to 20°C (Figure 3, Table S8).

### Warm versus cold thermal limits and seasonally acclimatized fish

A group of carnivorous fish were tested after an extended acclimation period to 20°C (2 and 4 weeks) (Figure S1). Peak *f*_Hmax_ increased within this timeframe while T_PEAK_ and T_ARR_ did not significantly change, indicating the opaleye could have continued acclimating a bit more if the acclimation time had been extended to 4-weeks across treatments (Figure S1). The fish tested in the acclimation rate experiment had an average peak *f*_Hmax_ of 207.78 ± 6.04 and 31.4 ± 0.3°C T_ARR_ after 14 days at 20°C compared to 213.20 ± 5.94 bpm and 32.7 ± 0.4°C from opaleye held 4 weeks at 20°C. Thus, the majority of their acclimation response was complete by week 2 post-temperature change, which is a more representative timescale for what they experience in the wild.

On the last timepoint (day 14 at 20°C), both upper and lower thermal limits for *f*_Hmax_ were assessed. Diet did not affect either upper (see Table S8) or lower thermal limits (Diet effect on T_ARR-cold_ χ^2^ = 0.660, df = 2, pr>χ^2^ = 0.719). For the cold ABT test, T_ARR - COLD_ was 5.3 ± 0.2°C for the herbivorous, 5.5 ± 0.5°C for the omnivorous, and 5.8 ± 0.6°C for the carnivorous.

Seasonally acclimatized opaleye (i.e., tested immediately following wild capture) experienced an average intertidal temperature of 18.4 ± 1.0°C (mean ± SD) in the 2 weeks prior to field sampling. Fish sampled in the late summer (August) had cardiac and morphometric performance that was closest to the herbivorous treatment at 20°C (Figure 4), with an average peak *f*_Hmax_ of 185.96 ± 7.29 bpm and T_AB_ of 23.4 ±0.3°C, T_PEAK_ of 28.9 ± 0.6°C, and T_ARR_ of 30.6 ± 0.7°C. Further, hepatosomatic index (HSI) was an average of 1.1% and relative ventricular mass (RVM) was 0.059%, where HSI was most similar to the herbivorous treatment at 12 and 20°C (1.1 and 0.8%, respectively).

**Figure 4.**
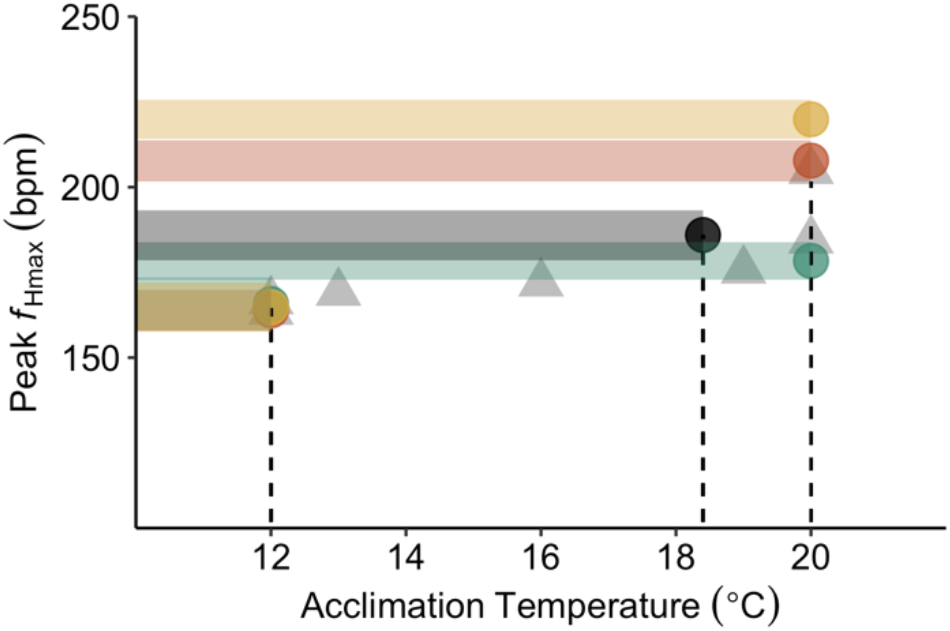
Comparison of treatment to opaleye literature values in peak *f*_Hmax_. Comparing mean peak *f*_Hmax_ values from current treatments to seasonally acclimatized opaleye as well as all other available literature values on opaleye [28,40]. Light Gray filled triangles indicate mean values from other studies on opaleye. Colors distinguish diet treatments used in this study (red: carnivorous, green: herbivorous, yellow: omnivorous). Dark gray circle indicates mean seasonally acclimatized opaleye response. Shaded rectangles indicate ± SEM for each treatment in this study. Dashed lines indicate acclimation or acclimatization temperature (from prior 2 weeks) in this study.

### Cardiac Fatty Acid Analysis

Diets and ventricles differed in their fatty acid composition (Table S4, Table S5, Figure S2). The algae had lower fatty acids overall and displayed higher proportions of saturated fats compared to brine shrimp (Table 4). In the ventricle, however, the DBI was highest in the 20°C herbivorous treatment, which was largely driven by differences in DHA content at that time (Table S5, Figure S2). Fish in the herbivorous treatment also had lower amounts of fatty acids with 18 carbon chains (e.g., oleic, linoleic, γ-linolenic) compared to the carnivorous and omnivorous treatments (Table S5, Figure S2).

Variation in *f*_Hmax_ could be partially explained by differences in ventricle DBI. For example, in 20°C acclimated fish, there were significant positive relationships between ventricle DBI vs. peak *f*_Hmax_ (p = 0.002; R^2^ = 0.32; Figure 5) and ventricle DBI vs. *f*_Hmax_ at 20°C (p = 0.009; R^2^ = 0.24; Figure 5). However, there was no relationship between DBI vs. peak *f*_Hmax_ in 12°C acclimated fish (p = 0.182; R^2^ = 0.07; Figure 5). In contrast, 12°C acclimated fish showed a significant positive relationship between DBI vs. *f*_Hmax_ at 12°C (p = 0.018; R^2^ = 0.25; Figure 5). DBI and average chain length were highly positively correlated, irrespective of diet (p < 0.001; R^2^ = 0.98). Thus, similar trends in average chain length vs cardiac parameters were observed. Several other significant correlations existed between cardiac parameters and specific fatty acid content (Figure 5d; Figure S3).

**Figure 5.**
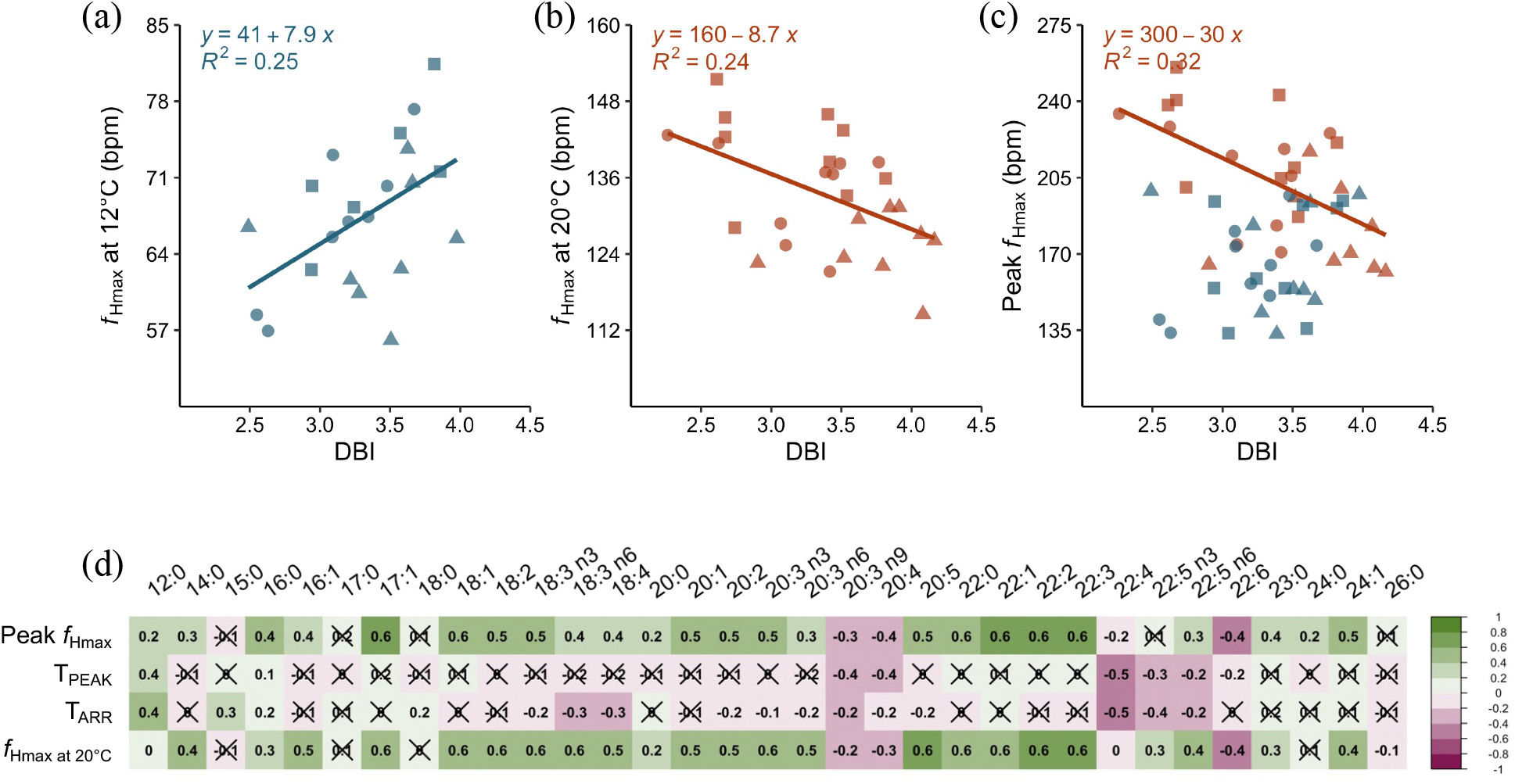
Fatty acid analysis from opaleye ventricles. Plots a-c show linear regressions between DBI and *f*_Hmax_ from opaleye across all diet treatments, where shapes indicate different diet treatments (carnivorous = circle, herbivorous = triangle, omnivorous = square). a) *f*_Hmax_ at 12°C from 12°C acclimated opaleye b) *f*_Hmax_ at 20°C from 20°C acclimated opaleye c) Absolute peak *f*_Hmax_ from 12°C (blue) and 20°C (red) acclimated opaleye. Equations are presented for significant linear regressions between parameters (p<0.05). There was no significant relationship between peak *f*_Hmax_ and DBI for 12°C acclimated fish. d) Correlations between cardiac parameters and individual fatty acids or fatty acid summary metrics from 20°C acclimated fish ventricles (made using ‘Corrplot’ verson 0.84 in R). Numbers and colors indicate the correlation coefficient. Non-significant correlations are crossed out (alpha < 0.05).

## Discussion

Diet is an understudied factor that could impact the rate and capacity for thermal acclimation. Here, we measured timescales of cardiac remodeling in a temperate omnivorous fish, opaleye, during a simulated warming event to determine whether differences in the opaleye’s diet influenced the rate of remodeling or overall capacity for thermal plasticity. We found that diet does influence the acclimation capacity for *f*_Hmax_ but not the rate of acclimation. We also examined how cardiac thermal performance related to ventricle fatty acid composition and found significant relationships between fatty acid composition and *f*_Hmax_. This has important implications for our understanding of species vulnerability to rising temperatures.

### Thermal acclimation capacity was diet dependent

In the face of ongoing anthropogenic changes to environmental temperatures and food resources [7], it is critical to consider if nutrition can mediate ectotherm resilience. Here, diet clearly impacted cardiac remodeling for *f*_Hmax_, but it did not alter the acclimation capacity of thermal limits (T_AB_, T_PEAK_, or T_ARR_). In all diet treatments, thermal limits increased with warm temperature acclimation, consistent with cardiac responses observed in opaleye and many other fishes [12,28]. For this generalist omnivorous species, a short-term shift to a lower energy, lipid, and protein diet did not constrain the thermal limits or the ability to acclimate them to a higher temperature. Further, the more nutritionally diverse (omnivorous) diet treatment here did not result in higher thermal limits compared to their carnivorous counterpart.

Instead, the height of the acute TPC for *f*_Hmax_, but not the breadth of the TPC, was sensitive to diet. Consequently, diet determined thermal plasticity for *f*_Hmax_, with the herbivorous treatment undergoing the lowest change in peak *f*_Hmax_ between the 12 and 20°C acclimated fish. Similar diet effects on TPCs for *f*_Hmax_ have been found in two other studies [28,47]. In Hardison *et al*. [28], *Ulva* sp. consumption lowered peak *f*_Hmax_, but not T_AB_, T_PEAK_, or T_ARR_ in opaleye at 20°C (Figure 4). In Papadopoulou *et al*. [47], allicin supplementation reduced *f*_Hmax_ across temperatures in exercise trained trout. Our data add to growing evidence that the amplitude of the acute TPC is more plastic than the breadth of the curve. While our results are from measures of cardiac performance, other performance metrics may follow similar patterns. For example, in some cases aerobic capacity may be more sensitive in amplitude, rather than breadth [48,49]. Differences in the amplitude of acute performance curves can alter functional thermal tolerance. Even though thermal limits were not impaired by diet here, the reduced *f*_Hmax_ in the herbivorous treatment indicates that *f*_Hscope_ (max – resting heart rate) could have a narrower thermal breadth if there were not corresponding decreases in resting heart rate [50]. In this case, performance under realistic temperature scenarios, or functional thermal tolerance, may be impaired, while more commonly measured extreme thermal limits (e.g., CT_max_, LT_50_, T_ARR_) remain the same.

At least in the short term, algae consumption in this study was not associated with any decrease in cardiac thermal performance at 12°C. Notably, opaleye in the herbivorous treatment had lower HSI (1.1%) than the carnivorous (3.1%) and omnivorous (3.0%) treatments at 12°C, indicative of an energetic disadvantage. However, more research is needed to determine the energetic and performance costs associated with herbivory in active fishes at cold temperatures and how the ecological role of herbivorous and omnivorous fish will change with an increasingly warm and more variable thermal environment.

### Thermal acclimation rate did not change with diet

In temperate environments, temperature often changes on shorter timescales than it takes for full thermal acclimation to occur [5]. For the endemic ectotherms, plasticity within the first few days after a temperature shift may be more important than overall acclimation capacity. However, most thermal performance data is collected on ectotherms that are fully acclimated to treatment conditions (generally ≥ 2 weeks), leading to overestimations of real-world thermal limits and performance. Further, if diet can alter the rate of acclimation, this could incentivize temperate fish to modify their diet with temperature.

As expected, thermal limits increased throughout the 2-week acclimation period, but surprisingly, there were no differences between diets. We did not observe any differences in acclimation rate despite HSI being higher in the carnivorous and omnivorous treatments (indicative of greater energy stores) after the initial 2-week acclimation to 12°C. All thermal limits showed a similar logarithmic relationship with time, consistent with acclimation rate studies using a critical thermal maxima test to estimate thermal limits [51,52]. These results add further evidence that consuming a lower quality diet (i.e., algae) did not impair thermal limits in opaleye, at least on timescales that are environmentally relevant.

Thermal limits were not affected by diet, but peak *f*_Hmax_ was, where fish from early timepoints had similar cardiac performance irrespective of diet, but performance diverged between the herbivorous and carnivorous/omnivorous treatments during the last week of acclimation to 20°C. Given the greater nutritional diversity (e.g., availability of micro and macronutrients), and flexibility in the omnivorous treatment and that opaleye, and many other aquatic omnivores [21], appear to consume greater proportions of algae in their diet in warmer water, we suspected the fish in the omnivorous treatment would have faster acclimation rates relative to those in the carnivorous and herbivorous treatments. However, we did not observe this. While fish in the omnivorous treatment had higher *f*_Hmax_ across temperatures than those in the herbivorous treatment after 2 weeks at 20°C, they had similar performance to the carnivorous treatment across all metrics and timepoints.

Notably, the fish at 12°C were much less likely to have food in their stomachs at the time of sampling, so it was not possible for us to estimate the exact ratio of algae to brine consumption at that time. However, we only started detecting small amounts of algae consumption on day 3 after raising the tank temperature to 20°C. The lack of differences between the carnivorous and omnivorous treatments may have resulted from fish not eating algae long enough for it to have a significant impact on their heart rate. While not within the scope of this study, it would be interesting to extend out to longer timepoints and see if prolonged acclimation to the omnivorous treatment at 20°C results in even greater algal consumption and higher cardiac performance. In Hardison *et al*. [28] opaleye acclimated to three weeks on an omnivorous diet (consisting of *Ulva* sp and brine shrimp) had slightly reduced cardiac thermal performance relative to a carnivorous treatment, indicating that 3-4 weeks at a given temperature may be necessary to observe subtle diet effects. Consistent with this, Sandblom *et al*. [9] demonstrated that the metabolism during digestion (or specific dynamic action) and standard metabolic rate took between 4 to 8 weeks to fully acclimate after switching from 10 to 16°C in shorthorn sculpin (*Myoxocephalus scorpius*).

While this is the first study that we are aware of to measure diet effects on thermal acclimation rates, there are a small number of studies that have measured how quickly cardiac performance acclimates in fish [13,18,53]. Interestingly, all studies found that temperate fish rapidly acclimate, with the majority of their response occurring within 3 days. Ekstrom *et al*. [13] measured *f*_Hscope_ in rainbow trout (*Oncorhynchus mykiss*) acclimated from 9 to 16°C and tested at 16°C. *f*_Hmax_ rapidly increased after 1 day at 16°C, and then began decreasing as acclimation occurred. These results were consistent with data from the herbivorous treatment, where *f*_Hmax_ at 20°C increased within 3 days of acclimation and then went back down by 2-weeks at 20°C. In contrast, the *f*_Hmax_ for the carnivorous and omnivorous treatments remained high throughout acclimation to 20°C. Gilbert *et al*. [18] used the ABT test to determine how *f*_Hmax_ changed during acclimation from 10 to 18°C in rainbow trout. In line with our results here, trout also underwent >50% of the acclimation response for *f*_Hmax_ in the first 72 hours post temperature change.

The speed with which temperate fishes, like trout and opaleye, mount an acclimation response indicates that acclimation rate may be a conserved and selected upon factor, although this remains to be tested. This work demonstrates how essential it is that we integrate time of exposure into our modelling of thermal tolerance and predictions of species responses to climate change. While these fish rapidly acclimate, they are still most vulnerable in the first few days after a temperature change, as their thermal tolerance has not fully compensated, and they are actively remodeling. It is logistically challenging to gather these data, but once in hand, they can be used to predict performance in complex environments, under variable, more realistic conditions.

### Fatty acid composition

Fatty acids serve many important functions, including as the phospholipid tails in biological membranes. Changes in fatty acids and membrane performance can impact cellular processes occurring across them, such as membrane bound enzyme activities, membrane potential, and ion movement [1,30], which can have consequences for organ and whole-animal performance. Although membrane fatty acid composition is tightly regulated, composition and content differ greatly across diets and membrane composition can be related to the fatty acid composition of the animal [54,55]. Consequently, dietary fatty acid composition can influence cardiac function, whole-animal metabolism, and swimming performance in fish [34–35,53]. For example, in Chatelier *et al*. [55], seabass (*Dicentrarchus labrax*) had lower cardiac output and aerobic scope when fed a diet containing fish oil as opposed to canola oil or palm oil. The effect of dietary lipid on critical thermal maximum and lethal thermal limits has also been assessed in a handful of studies, with mixed results. Oil type did not influence upper thermal limits in juvenile angelfish [56], but total lipid composition (10 vs. 20%) had a significant effect on critical thermal maxima in juvenile barramundi [33]. The effect of dietary fats on cardiac thermal plasticity in fish is relatively unknown. In our study, ventricle fatty acid composition was related to *f*_Hmax_ and acclimation temperature.

DBI is a measure of how “unsaturated” a tissue sample is on average, with higher DBI indicating a greater presence of double bonds in the fatty acid tails [57]. Here, cold-acclimated fish had higher heart rates (improved cardiac performance) when their ventricle fatty acids had more double bonds (DBI), or more fluid membranes (thus counteracting the cold). The opposite trend was observed for warm-acclimated fish: they had improved cardiac performance when their ventricles had fewer double bonds, or reduced membrane fluidity. In particular, saturated fatty acid (SFA) content appeared to drive the relationship between *f*_Hmax_ and DBI in 12°C acclimated fish, while highly unsaturated fatty acids (HUFA) appeared to be the major driver of the negative relationship between DBI and *f*_Hmax_ /peak *f*_Hmax_ in 20°C acclimated fish. DBI and average chain length were also positively correlated, meaning that fish with more unsaturated fats in their ventricle also had longer FA chains. While adding double bonds lowers the melting point and increases membrane fluidity, lengthening the carbon chain raises the melting point and increases membrane stability [1]. Thus, the fish may have increased the carbon chain length to partially compensate for their higher DBI.

Even though the algal dietary fatty acid composition had a higher proportion of saturated fats, the herbivorous treatment at 20°C had the highest ventricle DBI, which was largely driven by differences in the HUFA, DHA. DHA is an essential fatty acid critical to cognitive function and development [58]. However, it has six double bonds, resulting in a melting point of −44°C. It is unclear why the herbivorous fish ventricles contained more HUFAs, like DHA, relative to SFAs at 20°C. One potential explanation is that the herbivorous fish were metabolizing the SFAs or prioritizing their deposition in other tissues, as SFA have higher energy density compared to HUFAs of the same length (although digestibility is lower). Given the lower lipid content of the herbivorous diet, the fish could have prioritized meeting their metabolic demand over optimizing their membrane fluidity.

Given the high degree to which cells can regulate membrane performance and the many other functions for lipids in the body, it is not surprising that only ~20-30% of the variation in *f*_Hmax_ could be explained by fatty acid composition. Christen *et al*. [59] found significant negative linear relationships between thermal limits (time to CT_max_) and omega-3 fatty acid composition in *Salvelinus alpinus, Salvelinus fontinalis*, and their hybrids. Here, there were some significant correlations between specific fatty acids and cardiac thermal limits, although there was no significant linear regression found between overall DBI and cardiac thermal limits. While not evaluated here, other factors, such as sterol composition [60], phospholipid class [61] and phospholipid fatty acid composition [62], are known influencers of membrane performance that may have differed between the fish ventricles at the time of sampling and could explain more of the variation in heart rate across treatments.

The global availability of the essential omega-3 fatty acids, DHA and EPA, are expected to decrease with increasing temperatures associated with climate change [31]. While there is concern over the predicted reduction in the global availability of omega-3 FAs, the results here indicate that higher tissue SFA is associated with improved cardiac performance in warm for opaleye. In other words, shifts in global FA composition may be of concern for human nutrition and other measures of fish performance (e.g., fish cognitive health), but could also mediate cardiac upper thermal tolerance in fishes.

## Conclusion

This study is the first we are aware of to assess the impact of diet on thermal acclimation rates in any ectotherm. While diet did not affect thermal limits for *f*_Hmax_ or acclimation rate, it did impact the amplitude of the acute thermal performance curve. This indicates that for generalist species, like opaleye, variation in diet is an important modulator of thermal plasticity on environmentally relevant timescales. During warming events, thermal performance changes as a function of exposure time, and TPC amplitude is dependent upon nutrition. Mechanistically, fatty acids were related to heart performance in ways predicted by homoviscous adaptation, indicating that fatty acids may be a primary regulator of cardiac thermal performance in ectotherms. While temperate fish, like opaleye, have a remarkable capacity for rapid thermal plasticity, they are still most vulnerable in the first few days after a warming event, as they have yet to fully compensate for changes in temperature. These results highlight the importance of accounting for changes in nutrition, thermal exposure time, and animal diet preference when assessing ectotherm thermal tolerance.

## Acknowledgements

We thank Lucy Johnson, Cam Blair, Andrea Chandler, Bella Giglio, and Yvette Gaytan for assistance in the lab and with fishing; Terra Dressler, Jasmine Childress, Krista Kraskura, Jacey Van Wert, Claire Gonzalez, Jessica Madden, Bashir Ali, Hope Hardison for help fishing; David Davis, Christoph Pierre, and the entire marine operations staff at the University of California, Santa Barbara for making our science possible; Dr Elizabeth Wilbanks for lending equipment; and Dr Christopher Jerde, Dr Gretchen Hofmann, and Dr Elizabeth Wilbanks for advice and feedback.

