## Supplementary Information for "Diet changes thermal acclimation capacity, but not acclimation rate in a marine ectotherm (*Girella nigricans*) during warming"

**Table S1.** Ibutton data from temperature acclimations

| Replicate | 12°C | | | 20°C | | |
|---|---|---|---|---|---|---|
| | **Carn** | **Herb** | **Omni** | **Carn** | **Herb** | **Omni** |
| 1 | 12.1 ± 0.7 | 11.5 ± 0.5 | 12.6 ± 0.5 | 20.1 ± 0.7 | 19.6 ± 0.8 | 20.4 ± 0.5 |
| 2* | 12.3 ± 0.4 | 12.1 ± 0.6 | 11.7 ± 0.5 | 20.0 ± 0.5 | 20.6 ± 0.6 | 20.4 ± 0.5 |
| 3 | 12.0 ± 0.3 | 12.5 ± 0.3 | 12.1 ± 0.3 | 19.5 ± 0.3 | 19.9 ± 0.4 | 19.6 ± 0.5 |
| 4 | 11.6 ± 0.2 | 11.9 ± 0.2 | 12.5 ± 0.3 | 20.1 ± 0.2 | 20.0 ± 0.3 | 20.8 ± 0.3 |
| 5 | 12.2 ± 0.4 | 12.6 ± 0.2 | 12.3 ± 0.4 | 20.2 ± 0.3 | 20.8 ± 0.2 | 20.7 ± 0.3 |

Represented are mean (±SD) for Ibutton (temperature logger, Maxim Integrated) data from each experimental replicate. *indicates some Ibutton data missing in middle, however daily checks were consistently within the temperature range.

**Table S2.** Stomach content information for omnivorous treatment

| Timepoint | # Fish with ogo detected | # Fish with brine shrimp detected | Ogo stomach contents (% body mass) | Brine shrimp stomach contents (% body mass) | Ogo:total stomach contents (%) |
|---|---|---|---|---|---|
| Before change (12°C) | 0/11 | 4/11 | 0.0000 | 0.0631 | 0.0000 |
| Day 1 at 20°C | 0/12 | 2/12 | 0.0000 | 0.1529 | 0.0000 |
| Day 3 at 20°C | 2/12* | 3/12 | 0.0005 | 0.1576 | 0.3163 |
| Day 7 at 20°C | 1/8** | 5/8 | 0.0000 | 0.2133 | 0.0000 |
| Day 14 at 20°C | 11/22*** | 14/22 | 0.1005 | 0.3344 | 23.1088 |

Represented are the number of fish with ogo and brine shrimp detected in their stomach contents at the time of sampling. The amount of each diet item found in the stomach is calculated as the percent body mass of the individual fish and is presented as the average across all fish at that timepoint. *1 fish regurgitated ogo during test and no brine shrimp was detected in the tank. Therefore, not included in % body mass calculations. ** Ogo detected in gut, but stomach was empty. Therefore, not included in % body mass calculations. ***For 2 fish there was no brine shrimp left in tank at the time of sampling. These are included in % body mass calculations.

**Table S3.** Dietary and whole-body proximate composition (% wet weight)

| **Dietary Proximate Composition (% dry weight)** | | | |
|---|---|---|---|
| | *Gracilaria pacifica* | *Artemia* sp. | |
| % Protein | 20.73 ± 2.86 | 28.34 ± 1.41 | |
| % Lipid | 3.70 ± 0.72 | 11.03 ± 0.30 | |
| % Ash | 16.69 ± 3.84 | 12.38 ± 0.62 | |
| **Whole Body Proximate Composition (% dry weight)** | | | |
| | Carn | Herb | Omni |
| % Protein | 37.41 ± 4.27[a] | 35.57 ± 2.02[a] | 43.02 ± 2.29[a] |
| % Lipid | 18.57 ± 0.97[a] | 7.24 ± 0.91[b] | 20.74 ± 0.67[a] |
| % Ash | 14.76 ± 1.51[a] | 23.40 ± 2.45[b] | 11.91 ± 0.52[a] |

Represented are means and standard error values for dietary proximate composition in *Gracilaria pacifica* (n = 6 per measure), *Artemia* sp. (n = 3 per measure), and proximate body composition from whole opaleye (minus some dissected out organs; n = 7-8 per diet treatment) sampled on day 14 post temperature change to 20°C. Proximate body composition for opaleye were statistically analyzed using 1-way ANOVA and post-hoc Tukey HSD when appropriate. Letters indicate significant differences between treatment groups.

**Table S4.** Dietary fatty acid composition

| Summary Statistics for diet samples (mean ± SEM) | | |
|---|---|---|
| **Fatty Acid** | **Brine (pmol/mg dry tissue mass)** | **Ogo (pmol/mg dry tissue mass)** |
| 12:0 | 128.53 ± 12.73 | 63.93 ± 14.92 |
| 14:0 | 1657.06 ± 159.76 | 859.68 ± 186.96 |
| 15:0 | 210.11 ± 15.99 | 43.41 ± 12.12 |
| 16:0 | 4294.66 ± 544.33 | 5989.52 ± 1318.51 |
| 16:1 | 6075.68 ± 557.75 | 254.34 ± 106.97 |
| 17:0 | 146.19 ± 12.99 | 14.35 ± 3.68 |
| 17:1 | 267.59 ± 21.37 | 10.76 ± 4.14 |
| 18:0 | 1805.23 ± 307.00 | 407.58 ± 84.66 |
| 18:1 | 12260.74 ± 1554.57 | 952.27 ± 232.51 |
| 18:2 | 7902.38 ± 798.00 | 100.43 ± 39.05 |
| 18:3:n3 | 21160.92 ± 2028.76 | 151.75 ± 125.90 |
| 18:3:n6 | 417.99 ± 63.66 | 12.63 ± 3.27 |
| 18:4 | 10300.76 ± 795.33 | 60.88 ± 41.12 |
| 20:0 | 18.12 ± 2.53 | 0.36 ± 0.36 |
| 20:1 | 184.85 ± 19.19 | 10.21 ± 2.01 |
| 20:2 | 241.25 ± 24.54 | 42.26 ± 8.72 |
| 20:3:n3 | 630.41 ± 58.71 | 5.06 ± 3.70 |
| 20:3:n6 | 124.87 ± 14.10 | 43.28 ± 8.44 |
| 20:3:n9 | 80.91 ± 11.19 | 18.07 ± 3.41 |
| 20:4 | 1681.76 ± 203.53 | 439.00 ± 85.99 |
| 20:5 | 4746.66 ± 366.01 | 81.71 ± 36.17 |
| 22:0 | 11.37 ± 1.42 | 4.78 ± 1.31 |
| 22:1 | 6.20 ± 1.41 | 9.01 ± 1.14 |
| 22:2 | 0.93 ± 0.08 | 2.28 ± 0.68 |
| 22:3 | 6.89 ± 1.25 | 0.14 ± 0.14 |
| 22:4 | 18.01 ± 1.85 | 4.00 ± 0.63 |
| 22:5:n3 | 63.48 ± 6.59 | 1.16 ± 0.82 |
| 22:5:n6 | 117.7 ± 15.13 | 2.43 ± 1.27 |
| 22:6 | 837.4 ± 59.68 | 9.46 ± 7.94 |
| 23:0 | 1.45 ± 0.17 | 0.09 ± 0.09 |
| 24:0 | 3.89 ± 0.46 | 3.17 ± 1.10 |
| 24:1 | 0.48 ± 0.16 | 23.13 ± 6.23 |

| | | |
|---|---|---|
| 26:0 | 1.67 ± 0.31 | 0 ± 0 |
| average chain length | 17.88 ± 0.01 | 16.46 ± 0.04 |
| DBI | 1.83 ± 0.02 | 0.46 ± 0.03 |
| HUFA | 8308.10 ± 658.63 | 604.29 ± 95.47 |
| MUFA | 18795.54 ± 1974.30 | 1259.72 ± 344.41 |
| PUFA | 40024.24 ± 3217.62 | 370.22 ± 208.94 |
| SFA | 8278.27 ± 959.33 | 7386.85 ± 1609.52 |
| total | 75406.44 ± 6579.36 | 9621.30 ± 2077.09 |

Means ± SEM for each fatty acid from diet samples. Fatty acids are also summarized by average chain length, number of double bonds (DBI), and amount of saturated fatty acids (SFA; no double bonds), monounsaturated fatty acids (MUFA; one double bond), polyunsaturated fatty acids (PUFA; 2 double bonds at any chain length or greater than 2 double bonds and less than 20 carbons in the chain), and highly unsaturated fatty acids (HUFA; greater than or equal to 3 double bonds and 20 or more carbons in the chain length). Samples are standardized to dry weight and presented as pmol/mg of dry tissue mass. Fatty acids are reported as chain length: number of double bonds.

**Supplementary Table S5.** Ventricle fatty acid composition

| Summary Statistics for ventricle fatty acids (mean ± SEM) | | | | | | |
|---|---|---|---|---|---|---|
| **Fatty Acid** | **Carnivorous (pmol/mg tissue)** | | **Omnivorous (pmol/mg tissue)** | | **Herbivorous (pmol/mg tissue)** | |
| | **12** | **20** | **12** | **20** | **12** | **20** |
| 12:0 | 15.59 ± 2.27 | 18.41 ± 1.91 | 12.31 ± 1.79 | 15.34 ± 1.40 | 16.00 ± 2.09 | 16.39 ± 1.99 |
| 14:0 | 136.07 ± 18.47 | 141.63 ± 19.32 | 133.40 ± 17.64 | 145.18 ± 14.85 | 92.32 ± 3.95 | 106.24 ± 5.71 |
| 15:0 | 52.13 ± 6.00 | 44.32 ± 3.04 | 49.72 ± 6.73 | 46.66 ± 2.97 | 45.43 ± 4.71 | 47.96 ± 4.11 |
| 16:0 | 4578.38 ± 480.57 | 4683.44 ± 343.79 | 4329.96 ± 450.18 | 4674.41 ± 380.33 | 4645.31 ± 550.05 | 3973.04 ± 365.53 |
| 16:1 | 544.14 ± 109.41 | 434.74 ± 56.81 | 588.10 ± 117.04 | 437.28 ± 50.4 | 314.76 ± 34.22 | 267.19 ± 15.88 |
| 17:0 | 92.60 ± 11.92 | 80.37 ± 5.26 | 84.84 ± 9.88 | 89.62 ± 7.18 | 85.42 ± 11.87 | 75.72 ± 9.07 |
| 17:1 | 61.11 ± 8.88 | 55.40 ± 2.98 | 60.32 ± 11.48 | 62.98 ± 7.15 | 37.99 ± 4.7 | 30.08 ± 2.87 |
| 18:0 | 2539.88 ± 361.60 | 2335.91 ± 155.54 | 2131.24 ± 244.57 | 2457.98 ± 220.51 | 2389.31 ± 289.46 | 2352.46 ± 228.23 |
| 18:1 | 4529.38 ± 751.19 | 4372.17 ± 570.61 | 4612.46 ± 702.39 | 4961.47 ± 865.48 | 1782.38 ± 241.04 | 1289.18 ± 123.98 |
| 18:2 | 1010.21 ± 155.47 | 1088.89 ± 108.89 | 970.44 ± 128.08 | 1146.77 ± 147.78 | 409.98 ± 48.51 | 242.10 ± 18.15 |
| 18:3:n3 | 1442.08 ± 332.62 | 1470.50 ± 220.14 | 1734.98 ± 313.34 | 1616.48 ± 258.19 | 256.34 ± 28.55 | 154.56 ± 21.71 |
| 18:3:n6 | 21.82 ± 3.60 | 19.93 ± 2.60 | 23.71 ± 2.36 | 20.41 ± 2.04 | 9.50 ± 0.96 | 7.76 ± 0.57 |
| 18:4 | 108.43 ± 24.04 | 89.41 ± 10.65 | 133.33 ± 19.25 | 96.04 ± 16.34 | 27.93 ± 2.65 | 21.78 ± 4.16 |
| 20:0 | 22.13 ± 3.94 | 14.14 ± 2.65 | 20.71 ± 3.90 | 17.57 ± 2.03 | 15.00 ± 2.18 | 13.86 ± 2.09 |
| 20:1 | 204.10 ± 35.68 | 138.43 ± 14.88 | 200.21 ± 30.87 | 149.32 ± 16.86 | 98.97 ± 11.37 | 72.90 ± 6.96 |
| 20:2 | 138.10 ± 15.79 | 123.64 ± 5.41 | 130.59 ± 16.24 | 129.89 ± 10.26 | 90.48 ± 7.85 | 63.54 ± 4.8 |
| 20:3:n3 | 211.89 ± 28.03 | 245.16 ± 15.21 | 230.06 ± 31.46 | 248.74 ± 24.89 | 94.04 ± 7.87 | 61.83 ± 8.35 |

| | | | | | | |
|---|---|---|---|---|---|---|
| 20:3:n6 | 212.26 ± 20.51 | 190.42 ± 13.90 | 221.90 ± 19.06 | 186.93 ± 13.17 | 137.73 ± 13.76 | 96.56 ± 6.11 |
| 20:3:n9 | 233.14 ± 35.10 | 226.50 ± 34.14 | 288.76 ± 30.02 | 269.04 ± 31.03 | 190.77 ± 31.78 | 254.56 ± 27.85 |
| 20:4 | 4554.23 ± 497.59 | 4440.62 ± 499.04 | 5296.24 ± 417.69 | 5014.89 ± 419.83 | 4015.09 ± 563.10 | 5078.91 ± 394.71 |
| 20:5 | 1918.87 ± 230.56 | 2073.28 ± 155.61 | 2281.73 ± 274.3 | 1891.19 ± 157.76 | 1399.57 ± 187.62 | 785.19 ± 79.69 |
| 22:0 | 7.29 ± 1.77 | 4.78 ± 1.24 | 5.73 ± 1.36 | 5.70 ± 1.11 | 3.16 ± 0.62 | 1.69 ± 0.44 |
| 22:1 | 27.78 ± 6.43 | 15.24 ± 2.24 | 22.79 ± 4.6 | 16.59 ± 3.13 | 9.12 ± 1.36 | 6.11 ± 0.89 |
| 22:2 | 9.88 ± 1.52 | 8.49 ± 0.80 | 8.38 ± 1.45 | 9.90 ± 1.55 | 4.30 ± 0.60 | 2.71 ± 0.36 |
| 22:3 | 9.74 ± 1.62 | 9.10 ± 0.79 | 9.29 ± 1.78 | 10.13 ± 1.63 | 4.10 ± 0.59 | 2.16 ± 0.40 |
| 22:4 | 283.62 ± 20.30 | 307.36 ± 28.93 | 296.83 ± 41.22 | 295.28 ± 16.64 | 271.49 ± 22.31 | 288.57 ± 21.99 |
| 22:5:n3 | 1865.78 ± 202.43 | 1877.32 ± 174.26 | 1942.69 ± 240.84 | 1839.16 ± 74.25 | 1578.26 ± 159.58 | 1477.07 ± 168.74 |
| 22:5:n6 | 225.27 ± 10.94 | 292.53 ± 20.34 | 233.42 ± 12.50 | 289.44 ± 9.26 | 224.41 ± 27.72 | 216.40 ± 15.51 |
| 22:6 | 8576.93 ± 916.91 | 8820.27 ± 999.92 | 10192.78 ± 818.83 | 8706.84 ± 996.29 | 9112.59 ± 1073.22 | 12088.71 ± 1536.26 |
| 23:0 | 2.13 ± 0.38 | 1.27 ± 0.32 | 1.52 ± 0.30 | 1.86 ± 0.38 | 1.43 ± 0.36 | 1.22 ± 0.27 |
| 24:0 | 3.66 ± 0.55 | 2.22 ± 0.49 | 2.67 ± 0.58 | 3.23 ± 0.76 | 3.64 ± 1.22 | 2.31 ± 0.68 |
| 24:1 | 12.43 ± 1.82 | 11.26 ± 1.15 | 8.87 ± 1.31 | 13.83 ± 1.98 | 10.67 ± 1.88 | 9.20 ± 0.94 |
| 26:0 | 1.21 ± 0.41 | 0.80 ± 0.46 | 1.06 ± 0.56 | 1.51 ± 0.97 | 2.91 ± 1.85 | 1.19 ± 0.81 |
| Chain Length | 19.44 ± 0.10 | 19.44 ± 0.14 | 19.61 ± 0.11 | 19.44 ± 0.14 | 19.7 ± 0.12 | 20.04 ± 0.11 |
| DBI | 3.15 ± 0.12 | 3.17 ± 0.16 | 3.38 ± 0.12 | 3.15 ± 0.16 | 3.41 ± 0.14 | 3.77 ± 0.13 |
| SFA | 7451.07 ± 802.35 | 7327.30 ± 465.78 | 6773.17 ± 709.04 | 7459.06 ± 604.58 | 7299.94 ± 841.02 | 6592.08 ± 600.46 |
| MUFA | 5378.94 ± 903.34 | 5027.24 ± 634.32 | 5492.74 ± 846.69 | 5641.47 ± 934.43 | 2253.88 ± 277.05 | 1674.66 ± 144.16 |
| PUFA | 2730.52 ± 516.94 | 2800.87 ± 343.35 | 3001.43 ± 468.02 | 3019.49 ± 417.67 | 798.53 ± 72.62 | 492.44 ± 39.43 |

| HUFA | 18091.73 ± 1442.71 | 18482.56 ± 1541.71 | 20993.7 ± 1178.24 | 18751.66 ± 1210.11 | 17028.04 ± 1401.78 | 20349.94 ± 1756.79 |
|---|---|---|---|---|---|---|
| total | 33652.29 ± 3054.71 | 33637.99 ± 1264.95 | 36261.02 ± 2414.87 | 34871.69 ± 1209.08 | 27380.48 ± 2107.91 | 29109.08 ± 2045.39 |

Means ± SEM for each fatty acid from ventricle samples taken after 2 weeks at 12°C and 2 weeks at 20°C. Fatty acids are also summarized by average chain length, number of double bonds (DBI), and amount of saturated fatty acids (SFA; no double bonds), monounsaturated fatty acids (MUFA; one double bond), polyunsaturated fatty acids (PUFA; 2 double bonds at any chain length or greater than 2 double bonds and less than 20 carbons in the chain), and highly unsaturated fatty acids (HUFA; greater than or equal to 3 double bonds and 20 or more carbons in the chain length). Samples are presented as pmol/mg wet tissue mass. Fatty acids are reported as chain length: number of double bonds. Carnivorous = brine diet, omnivorous = ogo + brine diet, herbivorous = ogo diet.

**Table S6.** Summary Statistics for all metrics from the Arrhenius breakpoint test.

| Timepoint | Diet | n | Mean ± SEM | | | |
|---|---|---|---|---|---|---|
| | | | Peak $f_{Hmax}$ | $T_{AB}$ | $T_{PEAK}$ | $T_{ARR}$ |
| Before change (12°C) | Carn | 10 | 163.82 ± 6.03 | 21.2 ± 0.6 | 26.3 ± 0.5 | 28.2 ± 0.6 |
| | Herb | 10 | 165.74 ± 7.81 | 20.4 ± 0.8 | 26.9 ± 0.6 | 28.0 ± 0.5 |
| | Omni | 11 | 164.78 ± 7.21 | 21.0 ± 0.5 | 27.1 ± 0.7 | 28.5 ± 0.9 |
| Day 1 at 20°C | Carn | 9 | 181.36 ± 6.72 | 22.5 ± 0.7 | 27.1 ± 0.7 | 28.6 ± 0.7 |
| | Herb | 11 | 174.56 ± 5.71 | 23.2 ± 0.3 | 28.6 ± 0.8 | 29.7 ± 0.9 |
| | Omni | 10 | 177.34 ± 6.79 | 23.6 ± 0.5 | 27.4 ± 0.4 | 28.3 ± 0.6 |
| Day 3 at 20°C | Carn | 11 | 193.94 ± 7.18 | 23.5 ± 0.4 | 28.4 ± 0.5 | 30.2 ± 0.5 |
| | Herb | 9 | 184.56 ± 5.25 | 23.3 ± 0.7 | 28.5 ± 0.3 | 30.2 ± 0.3 |
| | Omni | 11 | 189.83 ± 5.17 | 23.8 ± 0.4 | 29.5 ± 0.6 | 30.6 ± 0.7 |
| Day 7 at 20°C | Carn | 8 | 195.00 ± 4.49 | 24.1 ± 0.2 | 28.2 ± 0.4 | 29.9 ± 0.5 |
| | Herb | 8 | 186.03 ± 7.87 | 24.1 ± 0.2 | 28.5 ± 0.6 | 32.1 ± 0.3 |
| | Omni | 7 | 205.73 ± 10.58 | 24.2 ± 0.6 | 28.8 ± 0.6 | 31.3 ± 1.1 |
| Day 14 at 20°C | Carn | 12 | 207.78 ± 6.04 | 24.5 ± 0.4 | 30.0 ± 0.4 | 31.4 ± 0.3 |
| | Herb | 11 | 178.36 ± 5.47 | 24.4 ± 0.3 | 30.1 ± 1.0 | 32.1 ± 0.9 |
| | Omni | 15 | 219.88 ± 5.65 | 25.4 ± 0.4 | 30.2 ± 0.4 | 31.9 ± 0.4 |

Means ± SEM for each test are presented. $T_{AB}$, breakpoint temperature of the heart; $T_{PEAK}$, temperature corresponding to maximum heart rate; $T_{ARR}$, temperature at the onset of cardiac arrhythmia; Peak $f_{Hmax}$, maximum heart rate achieved during the warm ABT test.

**Table S7.**

| SIC outputs for warm ABT test acclimation capacity polynomial curves | | | | |
|---|---|---|---|---|
| **Model #** | **df** | **SIC** | **ΔSIC** | **Formula** |
| Model 1 | 27 | 8848.30 | 0.00 | poly(temp, 3) * acc temp * diet + (1 \| fish_id) + (1 \| rep) |
| Model 2 | 33 | 8874.08 | 25.78 | poly(temp, 4) * acc temp * diet + (1 \| fish_id) + (1 \| rep) |
| Model 3 | 15 | 8966.85 | 118.55 | poly(temp, 4) * acc temp + diet + (1 \| fish_id) + (1 \| rep) |
| Model 4 | 13 | 8968.92 | 120.62 | poly(temp, 4) * acc temp + (1 \| fish_id) + (1 \| rep) |
| Model 5 | 13 | 8980.37 | 132.07 | poly(temp, 3) * acc temp + diet + (1 \| fish_id) + (1 \| rep) |
| Model 6 | 11 | 8982.16 | 133.86 | poly(temp, 3) * acc temp + (1 \| fish_id) + (1 \| rep) |
| Model 7 | 15 | 9172.63 | 324.33 | poly(temp, 3) * diet + (1 \| fish_id) + (1 \| rep) |
| Model 8 | 10 | 9175.00 | 326.70 | poly(temp, 3) + acc temp + diet + (1 \| fish_id) + (1 \| rep) |
| Model 9 | 11 | 9181.22 | 332.92 | poly(temp, 4) + acc temp + diet + (1 \| fish_id) + (1 \| rep) |
| Model 10 | 13 | 9185.91 | 337.61 | poly(temp, 4) + acc temp * diet + (1 \| fish_id) + (1 \| rep) |
| Model 11 | 7 | 9195.89 | 347.59 | poly(temp, 3) + (1 \| fish_id) + (1 \| rep) |
| Model 12 | 15 | 9602.05 | 753.75 | temp * acc temp * diet + (1 \| fish_id) + (1 \| rep) |
| Model 13 | 8 | 9633.44 | 785.14 | temp + acc temp + diet + (1 \| fish_id) + (1 \| rep) |
| Model 14 | 10 | 9634.98 | 786.68 | temp + acc temp * diet + (1 \| fish_id) + (1 \| rep) |
| Model 15 | 6 | 9636.42 | 788.11 | temp + acc temp + (1 \| fish_id) + (1 \| rep) |
| Model 16 | 5 | 9656.92 | 808.61 | temp + (1 \| fish_id) + (1 \| rep) |
| Model 17 | 7 | 9658.25 | 809.95 | temp + diet + (1 \| fish_id) + (1 \| rep) |

Represented are model formulas as input into R and SIC output results. df = degrees of freedom, SIC = Schwarz Information Criterion ΔSIC = SIC(model)—SIC(min SIC value), temp = acute temperature, acc temp = acclimation temperature (12 or 20°C), fish_id = individual fish., rep = replicate number of experiment.

**Table S8.** Statistics outputs from ANOVA for all metrics from the Arrhenius breakpoint test.

| Parameter | Factor | Chisq | Df | Pr(>Chisq) |
|---|---|---|---|---|
| Peak $f_{Hmax}$ | Timepoint | 71.6087 | 4 | <0.001 |
| | Diet | 14.3710 | 2 | <0.001 |
| | Diet x Timepoint | 19.4204 | 8 | 0.0128 |
| $T_{AB}$ | Timepoint | 120.9319 | 4 | <0.001 |
| | Diet | 3.6067 | 2 | 0.1647 |
| $T_{PEAK}$ | Timepoint | 57.8216 | 4 | <0.001 |
| | Diet | 2.9669 | 2 | 0.2269 |
| $T_{ARR}$ | Timepoint | 69.3436 | 4 | <0.001 |
| | Diet | 3.2205 | 2 | 0.1998 |

**Table S9.** % Acclimated for thermal limits across timepoints

| | $T_{AB}$ (% acclimated) | | | $T_{PEAK}$ (% acclimated) | | | $T_{ARR}$ (% acclimated) | | |
|---|---|---|---|---|---|---|---|---|---|
| **Diet** | **Day 1** | **Day 3** | **Day 7** | **Day 1** | **Day 3** | **Day 7** | **Day 1** | **Day 3** | **Day 7** |
| Carn | 39.4 | 69.7 | 90.9 | 21.6 | 56.8 | 51.4 | 12.5 | 62.5 | 53.1 |
| Herb | 68.3 | 70.7 | 80.5 | 53.1 | 50.0 | 50.0 | 41.5 | 53.7 | 100.0 |
| Omni | 59.6 | 63.8 | 74.5 | 9.7 | 77.4 | 54.8 | -5.9 | 61.8 | 82.4 |

Represented are the percent (%) of the total acclimation response for cardiac thermal limits (day 14 at 20°C - day 14 at 12°C) that was achieved on day 1, 3, 7 during acclimation to 20°C. Values are calculated as 100 x (thermal limit on day X at 20°C - thermal limit on day 14 at 12°C) / (thermal limit on day 14 at 20°C - thermal limit on day 14 at 12°C), where X = the day of testing. $T_{AB}$ = breakpoint temperature of the heart; $T_{PEAK}$ = temperature corresponding to maximum heart rate; $T_{ARR}$ = temperature at the onset of cardiac arrhythmia.

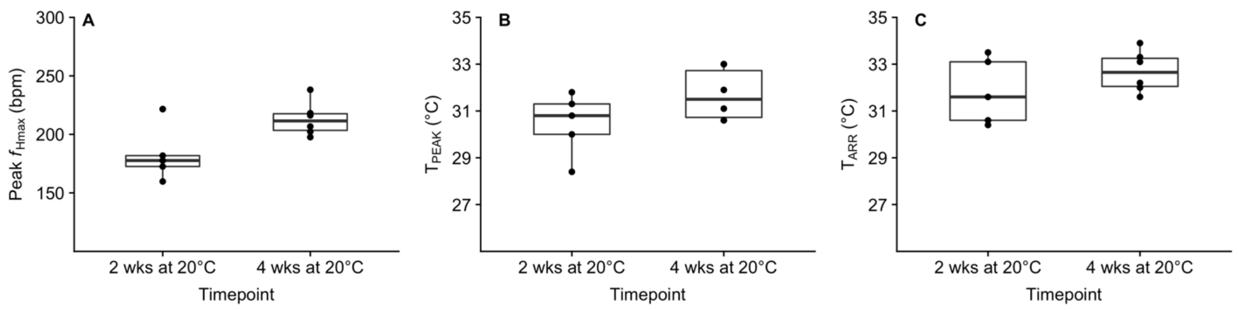


**Figure S1.** Cardiac performance in opaleye acclimated to 20°C for 2 or 4 weeks and fed a carnivorous diet. Presented are box plots of A) Peak $f_{Hmax}$ (bpm), B) $T_{PEAK}$ (°C) and C) $T_{ARR}$ (°C). Box plots represent interquartile ranges (boxes and whiskers), median values (solid lines) and outliers (> 1.5 beyond interquartile range) are plotted as data points outside the whiskers. Groups were compared using a two-tailed t-test. Peak $f_{Hmax}$ was found to be significantly different after 4 weeks compared to 2 weeks ($p = 0.026$). However, $T_{PEAK}$ and $T_{ARR}$ were not significantly different between timepoints ($p = 0.126$ and $p = 0.257$ respectively).

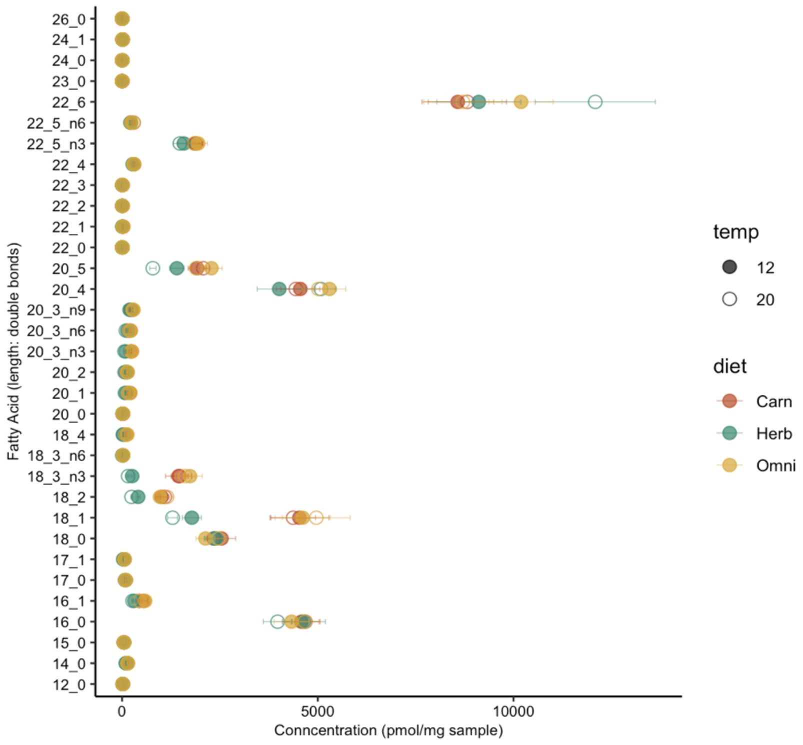


**Figure S2.** Mean ± SEM for various fatty acids in opaleye ventricles acclimated to 12°C (closed circles) and 20°C (closed circles). Colors indicate diet treatment, with carnivorous = red, herbivorous = green, omnivorous = yellow.

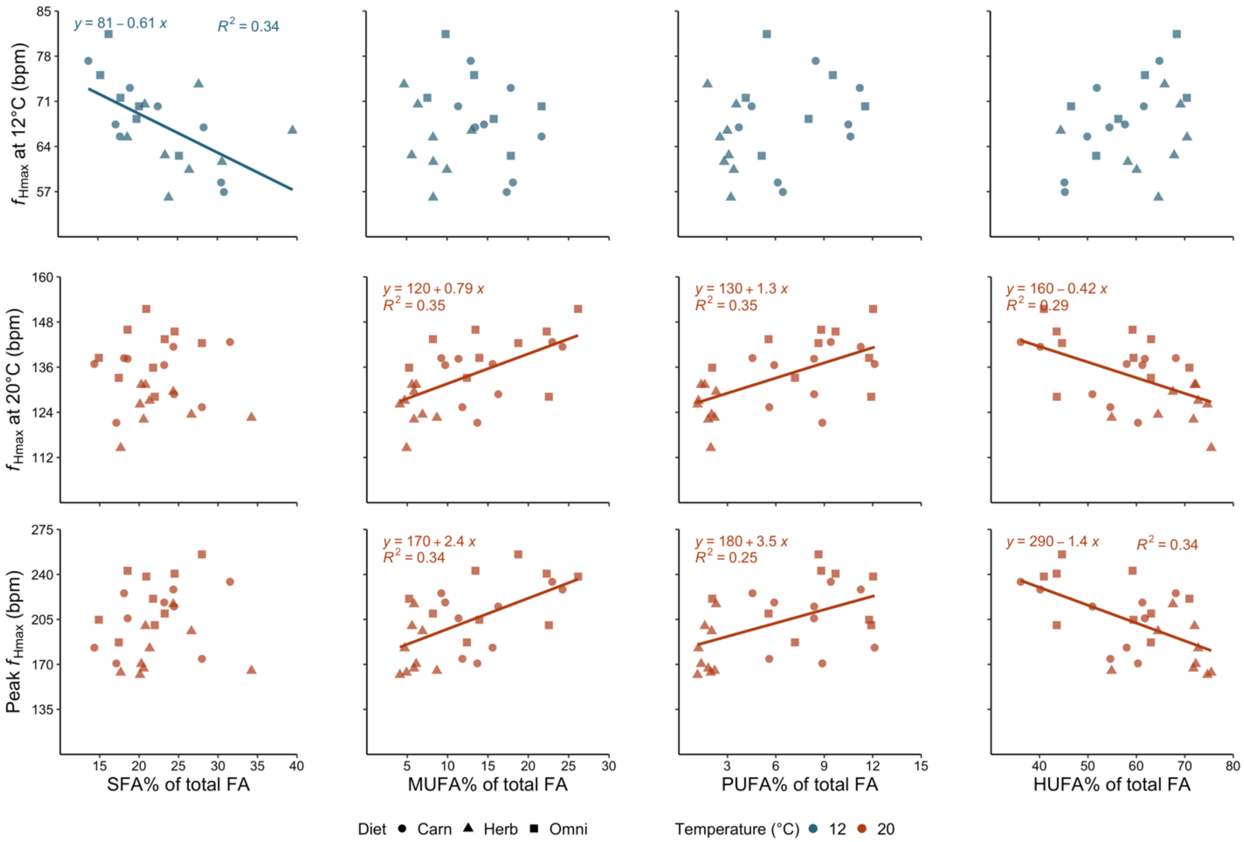


**Figure S3.** Correlations between various measures of max heart rate ($f_{Hmax}$) in 12 and 20°C acclimated opaleye and fatty acids. Each column is a different category of fatty acid presented as the proportion of the ventricle, with saturated fatty acids (SFA; no double bonds), monounsaturated fatty acids (MUFA; one double bond), polyunsaturated fatty acids (PUFA; 2 double bonds at any chain length or greater than 2 double bonds and less than 20 carbons in the chain), and highly unsaturated fatty acids (HUFA; greater than or equal to 3 double bonds and 20 or more carbons in the chain length). Lines are overlayed on plots where there was a significant linear regression between parameters (alpha <0.05). Equations and R2 values are provided in each significant plot. Colors indicate acclimation temperature (red = 20°C, blue = 12°C). Symbols indicate diets, with herbivorous = triangle, omnivorous = square, carnivorous = circle.
